# Microscale concert hall acoustics to produce uniform ultrasound stimulation for targeted sonogenetics in hsTRPA1-transfected cells

**DOI:** 10.1101/2021.08.21.457135

**Authors:** Aditya Vasan, Florian Allein, Marc Duque, Uri Magaram, Nicholas Boechler, Sreekanth H. Chalasani, James Friend

## Abstract

The field of ultrasound neuromodulation has rapidly developed over the past decade, a consequence of the discovery of strain-sensitive structures in the membrane and organelles of cells extending into the brain, heart, and other organs. Notably, clinical trials are underway for treating epilepsy using focused ultrasound to elicit an organized local electrical response. A key limitation to this approach is the formation of standing waves within the skull. In standing acoustic waves, the maximum ultrasound intensity spatially varies from near zero to double the mean in one half a wavelength, and can lead to localized tissue damage and disruption of normal brain function while attempting to evoke a broader response. This phenomenon also produces a large spatial variation in the actual ultrasound exposure in tissue, leading to heterogeneous results and challenges with interpreting these effects. One approach to overcome this limitation is presented herein: transducer-mounted diffusers that result in spatiotemporally incoherent ultrasound. The signal is numerically and experimentally quantified in an enclosed domain with and without the diffuser. Specifically, we show that adding the diffuser leads to a two-fold increase in ultrasound responsiveness of hsTRPA1 transfected HEK cells. Furthermore, we demonstrate the diffuser allow us to produce an uniform spatial distribution of pressure in the rodent skull. Collectively, we propose that our approach leads to a means to deliver uniform ultrasound into irregular cavities for sonogenetics.

## 1 Introduction

**S**ound diffusers have been applied to concert hall acoustics since the 1800s, when ornamentation along the walls or concave ceilings were used to introduce greater binaural dissimilarity^1^. Evolution of concert hall architecture eventually led to the development of the Schröder diffuser, arguably the first practical technique to disperse sound in a predictable manner. In the 1970s, Schröder proposed the phase grating diffuser^1,2^, a method to artificially create diffuse reflection. Composed of regular wells of different depths, these structures have periodicity in two dimensions as governed by a pseudostochastic sequence. In the typical configuration, waves incident on this structure undergo phase shifts corresponding to the depth of the wells through which they travel. The structure then scatters sound rather than reflecting it, depending on the magnitude of these phase shifts. This method has been widely adopted in architectural acoustics, where sound absorption—the only feasible alternative—is undesirable. This method has also been applied to ultrasound imaging^3^ and microparticle separation^4^ where sound absorption is likewise difficult. More recently, the principle of applying phase shifts to a coherent ultrasound field has led to development of acoustic holography^5,6^. This novel approach has enabled the generation of customized amplitude profiles based on the location and shape of the target region but does not enable the creation of spatiotemporally incoherent fields within an enclosed cavity.

Ultrasound transducers have been used for imaging tissue^7^, disrupting blood-brain barriers^8^, invasive^9^ and non-invasive neuromodulation^10^, and thrombolysis^11^. In these cases, ultrasound is typically focused at a certain depth defined either by the radius of curvature of the transducer, or an acoustic lens^12^, or phased arrays^13^. The fundamental limitation of these approaches is the formation of standing waves due to resonant reflections within the skull cavity formed by the relatively high impedance of the skull’s cortical bone compared to the tissue of the brain, and thus regions of either extremely high intensity or zero intensity at every one-half an acoustic wavelength^14^. The presence of these local maxima may lead to unintended bioeffects in tissues when applied to neuromodulation, including heating or even tissue damage. Such adverse effects in tissue have been reported during ultrasound-driven thrombolysis and blood brain barrier disruption^15,16^. Additionally, commonly used transducer materials such as lead zirconate titanate (PZT) also have limitations in high power applications at frequencies above ~ 1 MHz, producing losses, hysteresis, and internal (ohmic) heating as current passes through elemental lead present at the morphological grain boundary^17^. Another important consideration for sub-MHz frequencies, which have been used for neuromodulation studies in the past^18^, is the lowering of the cavitation threshold to a level that may elicit tissue damage at clinically relevant ultrasound amplitudes^19^. One approach to overcome these limitations is to build resonant devices using loss-free, single-crystal piezoelectric material operating in the 1–10 MHz range that are capable of delivering a spatiotemporally diffuse ultrasound field for various applications, including sonogenetics.

Sonogenetics relies on genetically engineering cells to be more sensitive to mechanical stimuli using membrane bound proteins^20,21^. This technique eliminates the need for focused ultrasound by ensuring that targeted neural circuits are the only ones that will respond to an ultrasound stimulus. Recent work has revealed that one protein in particular, human transient receptor potential A1 (*hsTRPA1*), produces ultrasound-evoked responses in several cell types^21^. One limitation of sonogenetics is that existing transducers producing planar or focused ultrasound are unsuitable. Furthermore, in many applications, the transducer must be small to avoid affecting animal behavior. Unfortunately, no small broadband transducers exist^22,23^ that might facilitate the generation of spatiotemporally random ultrasound noise from a similarly random input signal at sufficient power for sonogenetics. Moreover, commonly used animal models like rodents have small heads with a typical mass of 3-4g^24^, less than half the mass of all commercially available or research-based^25^ power ultrasound transducers known to the authors. The effective implementation of sonogenetics requires a very different transducer design. It must reduce interference between the radiated and reflected ultrasound, produce diffuse and uniform ultrasound throughout the region, transduce sufficient power to produce over 0.4 MPa acoustic pressure in tissue while remaining sufficiently small and light enough to attach to the head of a live mouse. In addition, these devices also have to avoid generating electromagnetic signals and localized temperature changes. If left to appear, these phenomena may conflate with the effects of ultrasound on the cells in sonogenetics experiments, reducing one’s confidence in ultrasound’s contribution to the observations. Transducers that can be attached to freely moving mice enable the study of neural circuits in their native state, without the confounding effects of anaesthesia as reported in past studies^26,27^.

We have overcome the limitations of existing transducers by incorporating a machined diffuser *on the transducer face* in order to produce spatiotemporally incoherent ultrasound. Diffusers are typically used in reducing coherent reflected sound—echoes—and their use *on the sound generator itself* has not been reported to the knowledge of the authors. A diffuser is ideally suited for sonogenetics as it nearly losslessly eliminates the presence of regions of either high or low intensity within an enclosed cavity, in both *in vitro* assays and within the rodent skull for longer-term applications. First, we discuss the design of the diffuser and validate its performance using numerical simulations. We then address the challenge of fabrication of a complex three-dimensional structure at sub-millimeter scales, as conventional photolithography, three-dimensional printing, and classic machining techniques are unsuitable for this task. Next, we characterize a device which has been coupled to a lithium niobate transducer operating in the thickness mode. We use lithium niobate, a single crystal material with low losses and no hysteresis over the MHz operating frequency range required for ultrasound neuromodulation^28^.

Forming uniform acoustic pressure fields in enclosed cavities is a key challenge in a variety of biomedical applications. Coherent propagation and reflection of ultrasound naturally forms antinodes and nodes in a cavity, corresponding to regions of strong and weak ultrasound. Uniform ultrasound reduces the risk of over or under-exposing the tissue regions that contain ultrasound-sensitive proteins. We present an application of the Schröder diffuser to screen for ultrasound-sensitive ion channels in human embryonic kidney (HEK293) cells *in vitro* for the purposes of identifying and isolating targets for sonogenetics in freely moving mice. We also verify the presence of nearly equivalent acoustic pressures across two deep brain regions in an *ex vivo* model.

## 2 Results

The design of the Schröder diffuser is based on quadratic-residue sequences defined by *s_n_* = *n*^2^, where *n*^2^ is the least non-negative remainder mod *N*, with *N* always an odd prime. One of the properties of this number sequence relevant to the design of an optimum diffuser is that both the Fourier transform of the exponential sequence *r_n_* = exp(*i*2*πs*_*n*_/*N*), and by extension the scattered wave produced by it, have a constant magnitude^1,29^ |*R_m_*| such that

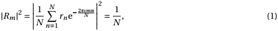

where 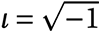.

We may then use this to define the wells’ depths, *d*(*x_n_, y_n_*), corresponding to the number sequence. In one dimension, the depth of the *n*^th^ well is given by^30^

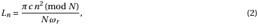

where *ω_r_* is the design frequency, *N* is a prime number, and *c* is the speed of sound in the medium. Extending the concept of a diffuser defined per the above numerical sequence to two dimensions involves replacing *n*^2^ in the above formula with *n*^2^ + *m*^2^, where *m* represents the number of wells in the second dimension. A representative image of a diffuser fabricated using a two-dimensional sequence is shown in Fig. 1.

**Figure 1:**
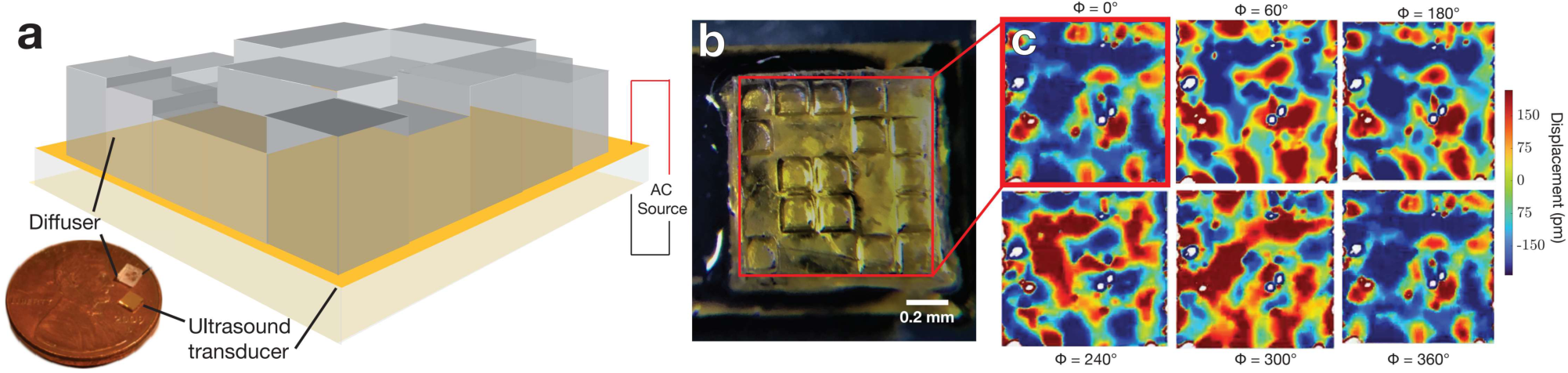
**(a)** A diffuser design based on Schröder’s method of quadratic-residue sequences to determine well depth. The wells were machined in glass using a KrF excimer laser system with a custom metal mask to restrict beam width. The machined depth of the pillars can be up to 309 *μ*m. **(b)** The glass diffuser block was then **(c)** bonded to a transducer operating in the thickness mode at 7 MHz using an ultraviolet light-curable epoxy. **(c)** A scanning laser Doppler vibrometer image of the diffuser face in the time domain shows phase differences corresponding to pillar heights (normalized autocorrelation > 0.73).

While a one-dimensional diffuser creates a uniform two-dimensional pressure field, a two-dimensional diffuser with varying well depths creates a uniform three-dimensional pressure field. Ultrasound neuromodulation typically relies on frequencies in the 1–10 MHz range^31^ and this requires sub-millimeter well depths as defined by eqn. (2). Although structures based on the quadratic-residue sequence have been achieved at the macro-scale in two dimensions and at the microscale in one dimension^4^, it has not been achieved in two-dimensional structures on the micron to sub-mm scale due to the lack of established fabrication techniques for these dimensions^32^. Conventional photolithography is good for creating patterns that have the same depth or, at most, a few different depths. It becomes challenging when features of varying depths are desired because multiple photolithography and etching steps are required. Alternate approaches, including three-dimensional or two-photon printing methods are unable to produce acoustically low-loss structures with sufficient dimensional accuracy at these scales. We sought to address these limitations by using an excimer laser to machine sub-millimeter pillars of varying heights in glass in two dimensions.

A time-domain scan using laser Doppler vibrometry (LDV; *see* Methods) in Fig. 1 shows significant phase correlation (normalized autocorrelation > 0.73) with the machined geometry. The transducer was driven at its resonance frequency with a sinusoidal input power range of 0.5 – 2 W and a peak pressure output of 0.6 MPa as measured with a fiber optic hydrophone^33^.

Finite element analysis (COMSOL 5.5, Comsol Inc., Los Angeles, CA USA) was used to validate the design of the diffuser. The domain was chosen to mimic an experimental setup used for identifying ultrasound sensitive ion channels in an *in vitro* setup. This consists of an inverted fluorescence microscope with a custom perfusion chamber to house a cover slip and transducer. The simulation domain is illustrated in Fig. 2 and specific dimensions of the domain and simulation parameters are included in the *Methods*. The transducer and the diffuser assembly was fixed at the bottom of the domain. A custom perfusion chamber that contains a slot for a coverslip was mounted over the ultrasound source. The transducer was coupled to the coverslip through water and there was a layer of media above the cover slip. The walls were defined to be hard boundaries with the acoustic impedance *Z*_i_ = ∞ such so that the normal derivative of the total acoustic pressure, 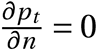.

**Figure 2:**
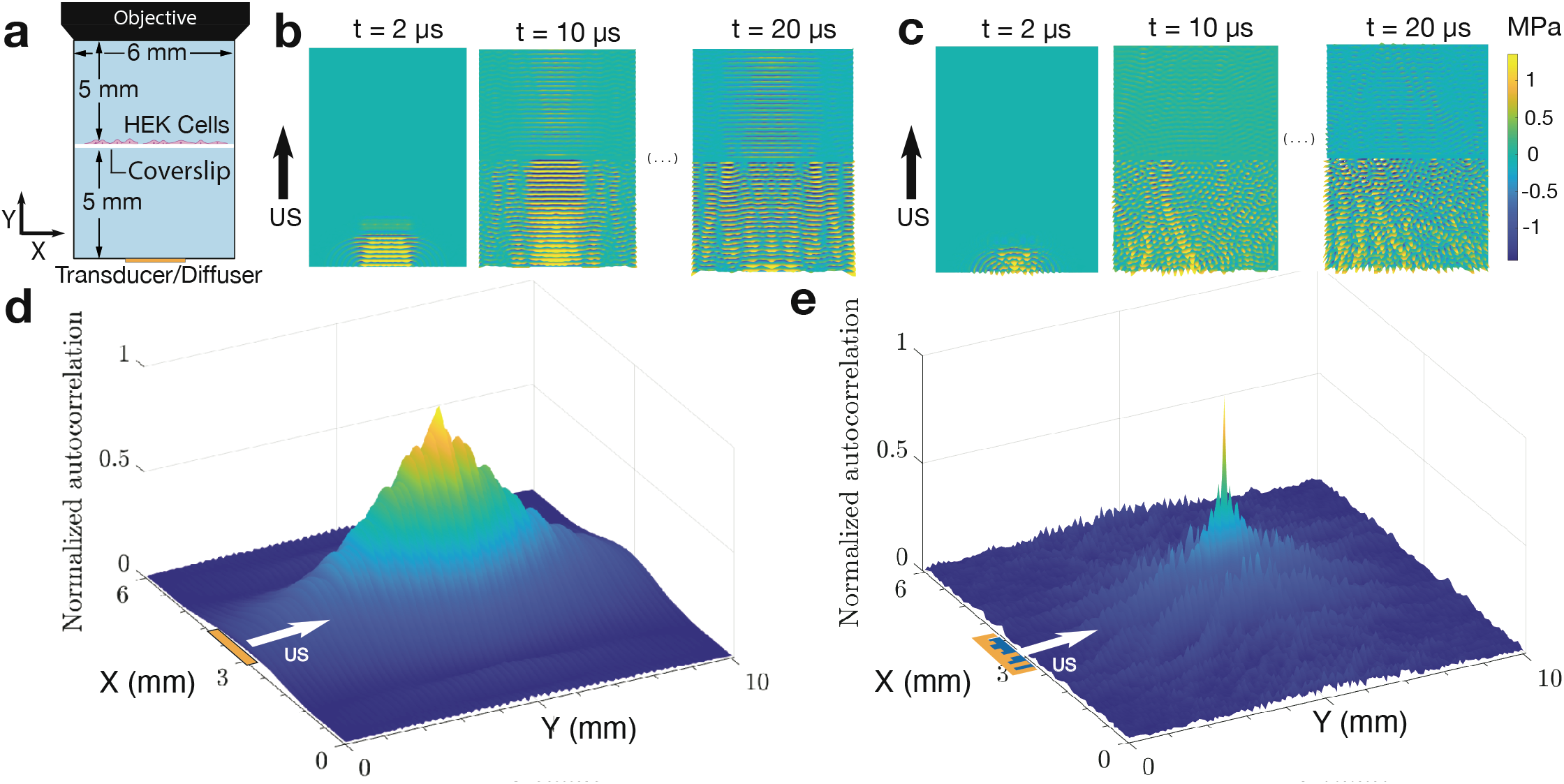
Two-dimensional pressure profile for the **(a)** domain **(b)** with and **(c)** with the diffuser. Human embryonic kidney (HEK) cells were placed in the middle of the (light blue) fluid domain with an objective lens for an inverted microscope at top. The pressure field was generated by defining a sinusoidal pressure displacement to the transducer face, located at the bottom of the domain. Pressure maps were extracted from the results in 1*μ*s time steps over grid points specified within the domain. A two-dimensional autocorrelation was performed on this grid over time; each X,Y point plotted from the **(d,e)** results of the autocorrelation corresponds to a single point in the (a) domain. Spatial and temporal periodicity are observed through the existence of a large value of autocorrelation over the domain **(d)** without the diffuser compared to the generally low autocorrelation with **(e)** the diffuser introduced onto the transducer.

The diffuser consists of seventeen elements, the heights of which were calculated from eqn. (2). The cover-slip in serves as a solid boundary and allows the evaluation of the acoustic field in the closed domain below and the open domain above it, corresponding to the different boundary conditions assigned to the model.

The time variation of the pressure field with and without the diffuser was evaluated (supplementary video S1 and S2). Several points in the fluid domain were chosen and the time evolution of the pressure field for the two cases was compared using the techniques described in the *Methods* section. A two-dimensional autocorrelation was calculated in order to determine if there were any locations within the domain with coherence (echoes) or localized increases or decreases (constructive and destructive interference) in ultrasound intensity.

Spatial and temporal patterns that form over the duration of the stimulus are represented by a two-dimensional autocorrelation in Fig. 2. It is evident that there is both spatial and temporal periodicity with the transducer alone (Fig. 2a and supplementary video S1) that is greatly reduced when the diffuser is introduced (Fig. 2b and supplementary video S2). Videos of the sample autocorrelation in the domain over the stimulus duration are presented in the supplementary information (videos S3 and S4) and show that there is greater autocorrelation over the duration of the stimulus without the diffuser (video S3). This indicates that the ultrasound field with the diffuser is temporally inconsistent. The autocorrelation plot is an instructive technique to determine periodicity in time and space but does not quantify spatial dispersion in the domain at the driving frequency.

For the purpose of quantifying the dispersion at 7 MHz, an isofrequency contour plot is provided in Fig. 3(a) without and (b) with the diffuser. Without the diffuser, wave vectors are only present in the vicinity of *k*_x_ = 0, along the direction of propagation of the pressure wave in the medium: the Y axis. The angular spread is 20° on either side of the direction of propagation without the diffuser. Particularly, the majority of the wave can be seen to be propagating along the Y axis, with significant sidelobes immediately to the left and right and much smaller sidelobes slightly farther away. Including the diffuser produces wave vectors beyond the main direction of propagation (Fig. 3b), with significant components oriented along directions from the Y axis (along *k*_x_) to the X axis (along *k*_y_). The previously significant sidelobes remain significant, but are augmented by wave propagation beyond 45° in the XY plane. This indicates strong dispersion in the domain when including the diffuser.

**Figure 3:**
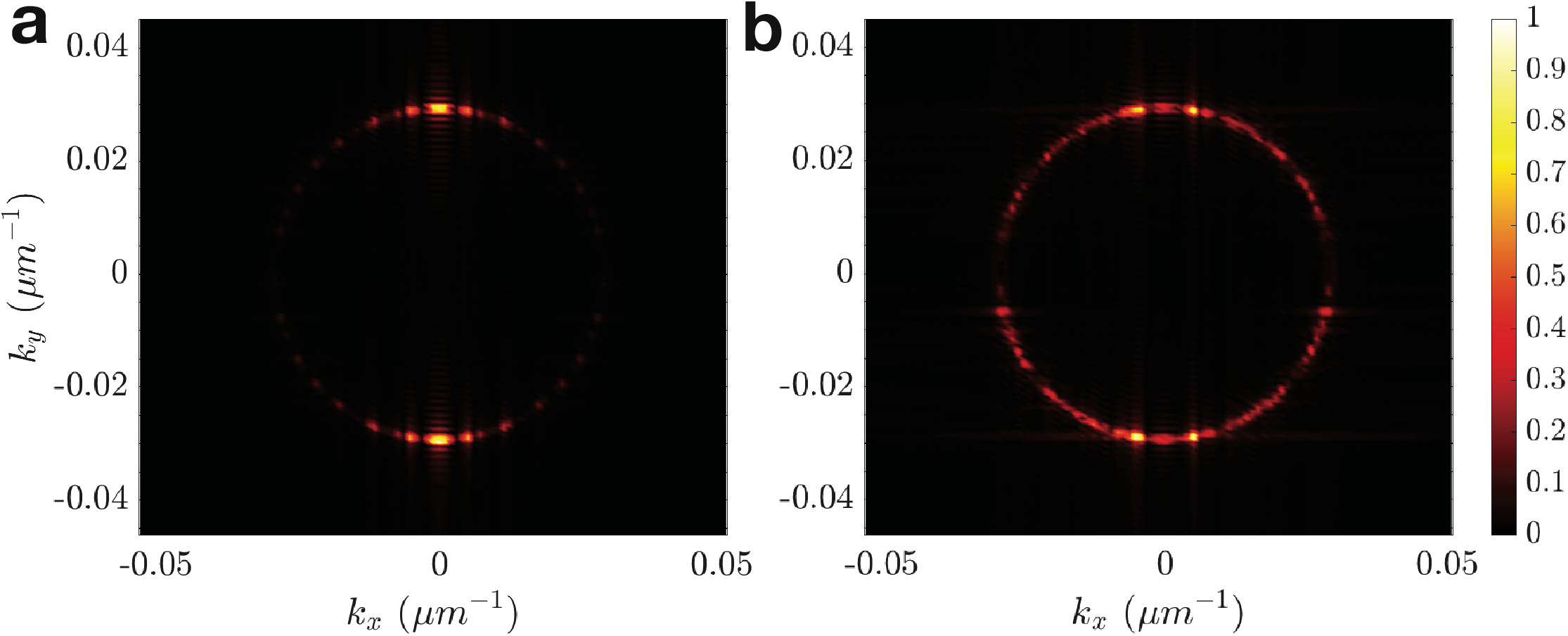
Isofrequency contour at the driving frequency **(a)** without and **(b)** with the diffuser. The circular profile traced by both cases corresponds to the wave vector in water at the driving frequency. The transducer (b) with diffuser produced wave vectors spread around this circular profile, indicating a more uniform distribution of the ultrasound. Without the diffuser, most of the wave is isolated to propagation along the Y axis.

To verify the effects of the diffuser *in vitro*, we used an upright optical imaging setup including an immersion objective, a custom perfusion chamber, and the diffuser assembly. The diffuser assembly and the test setup are represented in Fig. 4a; we used lithium niobate due to its relatively high coupling coefficient and zero hysteresis^34^ which implies no heating from the piezoelectric material itself. Human embryonic kidney (HEK293) cells expressing GCaMP6f^35^ were transfected with hsTRPA1. We compared fluorescence changes 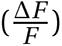 for four cases, with and without the channel, without the diffuser assembly (transducer alone), and with the diffuser assembly. Representative GCaMP6f images of HEK293 cells transfected with hsTRPA1 are shown in Fig. 4b and heat maps of fluorescence intensity with respect to time are presented in Fig. 4c, with a clear increase in both the magnitude and number of cells being activated with the diffuser assembly. Cells expressing hsTRPA1 and controls were tested at two different pressures, 0.32 MPa and 0.65 MPa. There was a consistent increase in fluorescence intensity with an increase in acoustic pressure for both the control and the hsTRPA1 condition, whether or not the diffuser was present. Including the diffuser increased the mean fluorescence amplitude by at least a factor of two for cells that had been infected with hsTRPA1 (*p* < 0.0001, Fig. 4d).

**Figure 4:**
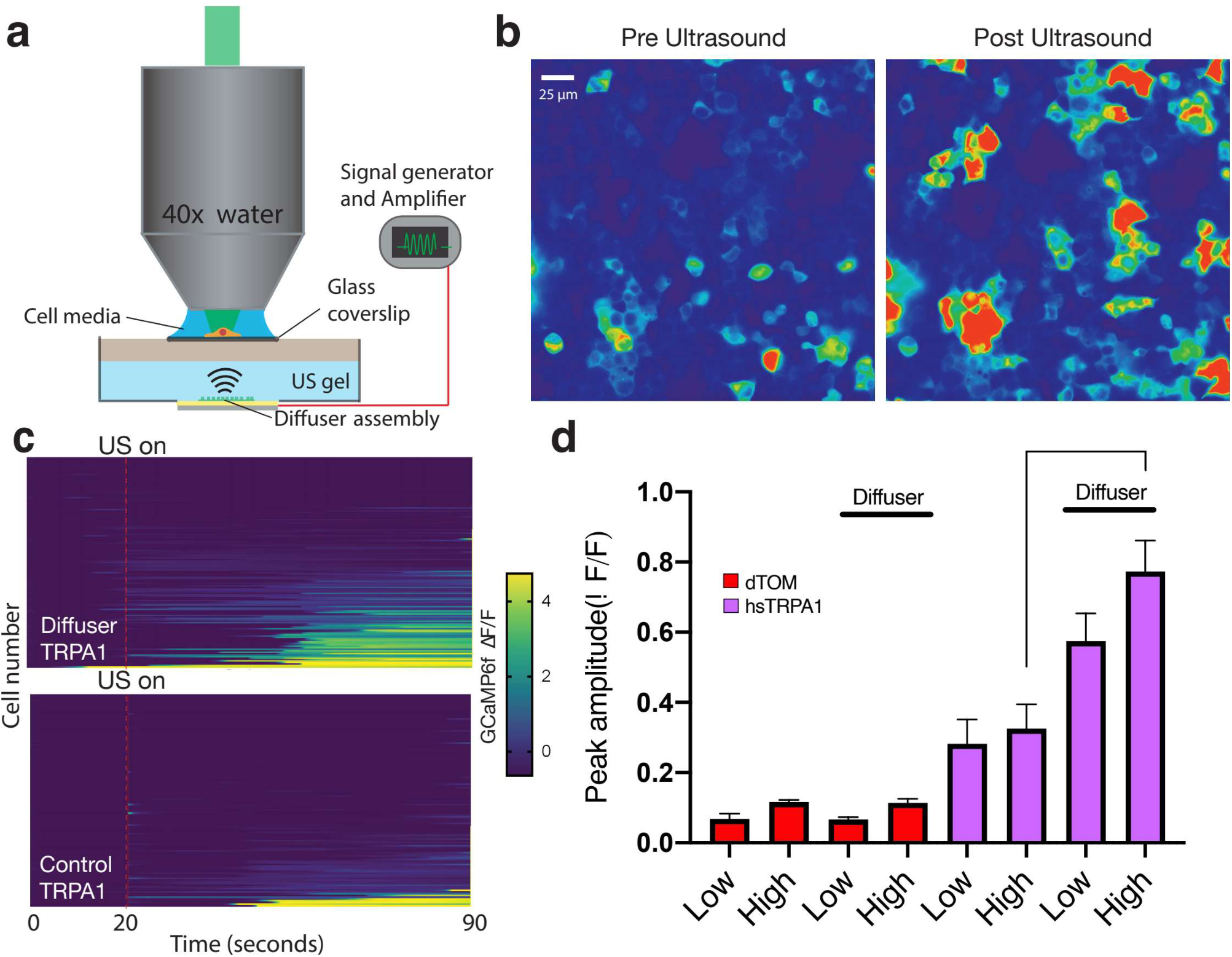
**(a)** The experimental setup for confirming the utility of the diffuser in an *in vitro* setting consists of a upright epi-fluorescent microscope, an immersion objective, and a chamber that houses cells on a coverslip and the diffuser assembly. Standing wave components may exist between the transducer and the cover slip and between the cover slip and the immersion objective. The calcium concentration before and after ultrasound stimulation in the same field of view is **(b)** shown for HEK cells expressing hsTRPA1. Comparison of fluorescence changes as measured using GCaMP6f reporters with respect to time for two cases, **(c)** without (control) and with the diffuser show an increase in both number and magnitude of cells being activated upon introduction of the diffuser. **(d)** HEK cells expressing TRPA1 show a greater response to ultrasound stimuli with a diffuser present in comparison to both no diffuser and dTom-based controls. The magnitude of the response when the diffuser is used is significantly greater (over twice as high) than when the diffuser is not used (*n* = 76−221, *p* < 0.0001 by a Mann-Whitney test).

We also tested the effects of ultrasound on mouse primary cortical neurons. Neurons were infected with adeno-associated viral (AAV) vectors to express hsTRPA1 and a genetically encoded calcium indicator, GCaMP6f^35^, or a control with only the calcium indicator. We found that ultrasound triggered an increase in calcium uptake in both cases, with the hsTRPA1 neurons showing a greater number of activated cells in comparison to the control (Fig. S2). A video of real-time calcium response in hsTRPA1-expressing neurons is presented in the supplementary information (Video S5).

The uniform nature of the ultrasound field created by the diffuser was also verified *ex vivo* in a mouse skull *see Methods*. Pressure measurements were taken at two different locations as indicated in Fig. 5 along the anterior-posterior axis, at the ventral surface of the pons and the ventral surface of the anterior olfactory bulb. With the diffuser, the pressure at both these locations was uniform, with minimal deviation between them and a uniform increase with input power to the transducer (Fig. 5). However, the transducer alone produced diverging values of pressure at these positions, so much so that the pressure at the pons (triangle) exceeded the pressure at the anterior olfactory bulb (circle) by a factor of 3 at an input power of 3 W, yet fell below the hydrophone’s minimum measurement value, 0.2 MPa, at the anterior olfactory bulb when using less than 1.25 W of power. By contrast, the diffuser had minimal deviation in pressure values at these locations, with pressure values ranging from 0.25 - 0.5 MPa at the ventral surface of both the pons and the anterior olfactory bulb. These brain regions were chosen not for their function, but as they were as they were the farthest from each other and the device, while keeping as much of the mouse skull intact during preparation [*see* methods]. Collectively, these results demonstrate that the diffuser is capable of delivering uniform ultrasound fields *in vivo* in comparison to the a transducer alone, thus enabling sonogenetic studies across large brain regions.

**Figure 5:**
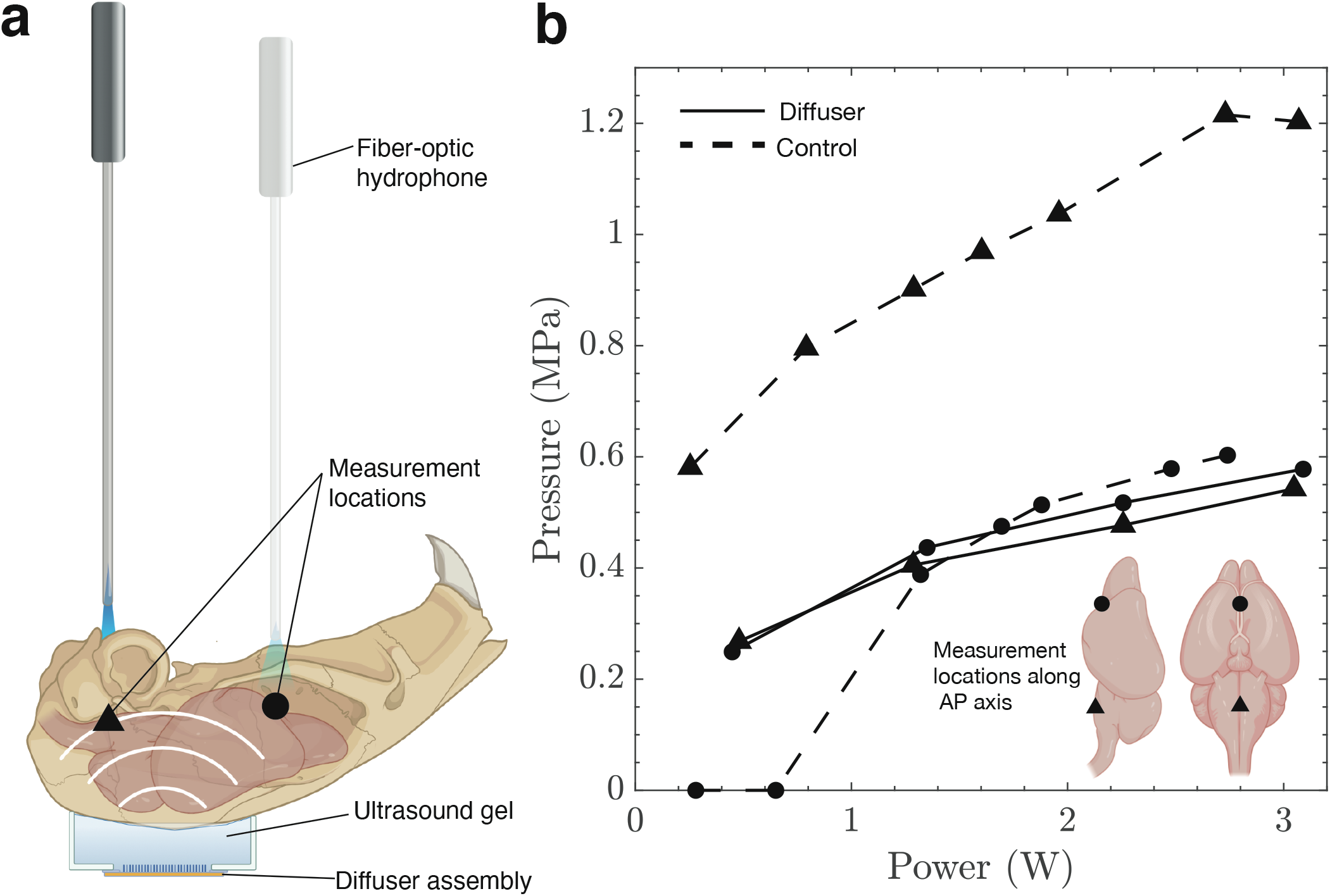
The pressure was measured using **(a)** a fiber optic hydrophone at two different locations along the anterior-posterior axis: the ventral surface of the pons (triangle) and the ventral surface of the anterior olfactory bulb (circle). The measured pressure is **(b)** uniform across different brains for different input powers above 0.2 MPa (minimum detectable pressure using our setup), indicating that the diffuser creates a uniform acoustic field within the skull cavity. This eliminates the influence of the cranial structure and ensures that only the brain regions that have been infected with hsTRPA1 will be sensitive to ultrasound stimuli. In comparison, the control case without the diffuser shows a three-fold deviation in pressure values for the same input power for different brain regions along the AP axis.

## 3 Discussion

Existing non- and minimally-invasive techniques to stimulate brain regions, such as transcranial magnetic stimulation and transcranial direct current stimulation, offer poor spatial resolution. This is a problem for precisely targeting brain regions that have specific functions. Ultrasound-based stimulation enables targeting brain regions with sub-millimeter-scale accuracy. This precision can be achieved in different ways, either by using an array to focus ultrasound to a specific region^36^ or by using sonogenetics to engineer cells to locally be more sensitive to mechanical stimuli.

Still, the limitation with focused ultrasound is the alteration in the position and shape of the focal zone due to spatial variations in acoustic impedance^37^. Sonogenetics is an attractive option because of the potential of having a toolkit of specific proteins that can be engineered to be sensitive to ultrasound stimuli at different frequencies or pressures. Current ultrasound transducers and how ultrasound interacts with the skull cavity are important limitations to translate Sonogeneticss. Standing waves in the skull cavity produce nodes and antinodes, each separated by one-half of the acoustic wavelength and responsible for pressure minima and maxima, respectively. This can lead to haemorrhage and heating in tissue^15^ in past studies. A further limitation is that ultrasound transducers cannot be driven using broadband white noise to produce a spatiotemporally random acoustic field at pressures sufficient to elicit cellular responses. The development of sonogenetics that started with the *TRP4* channel has expanded to include a library of proteins that are sensitive to ultrasound stimuli at different ultrasound stimulation parameters. Examples include MSC^38^, TREK^39^, Piezo^40^, and other TRP channels^41^, all have which have been shown to be sensitive to ultrasound *in vitro*.

Designed via computational analysis and fabricated with an excimer laser, a microscale Schröder diffuser was devised to eliminate the spatiotemporally heterogeneous distribution of ultrasound by placing it upon the emitting transducer. The transducer alone was shown to produce standing waves in the absence of the diffuser. With the diffuser in place, autocorrelation of the ultrasound field quantifies the elimination of the standing waves and consequent suppression of antinodes associated with potential tissue damage. This result is confirmed for the resonant frequency of operation of the transducer using an isofrequency contour. We verified the predictions of the simulation *in vitro* using HEK293 cells and neurons that were transfected with a sonogenetic candidate, hsTRPA1.

Development of sonogenetics in larger animal models—such as primates—will require ultrasound transducers that are capable of delivering an acoustic field that is spatially and temporally incoherent, as we have shown with the diffuser assembly. This ensures that the pressure in different brain regions is uniform over the stimulus duration, thus eliminating the aberrations in the acoustic field due to the skull cavity. Functionalization of specific brain regions using ultrasound-sensitive proteins can offer sub-millimeter spatial precision. Localization of Sonogenetic proteins in combination with an acoustic field provided by a diffuser assembly will ensure that the observed neuromodulatory effects are solely due to ultrasound activation of targeted regions of tissue and not due to the confounding effects of reflection or interference from the geometry of the skull.

## 4 Methods

### Ultrasound transducers

Ultrasound transducers used in this study were single crystal lithium niobate transducers operating in the thickness mode with lateral dimensions of 5×5 mm and thickness 500 *μ*m. The 128YX cut of lithium niobate was used and the fabrication process involved cleaning of the wafer with acetone, isopropyl alcohol, and ultra-pure deionized (DI) water followed by sputtering both sides with an adhesion layer of 20 nm titanium followed by 1 *μ*m gold. The deposition parameters were (with a Denton Discovery 635, Denton Vacuum LLC, New Jersey, USA) 5–10 nm of Ti at 1.2–1.6 A/s with the power set to 200 W, with argon as the gas in the chamber at 2.3 mT and the stage rotating at 13 RPM to ensure uniform deposition over the sample. The thickness of gold deposited was 1 *μ*m at a rate of 7 – 9 A/s.

### Diffuser fabrication and characterization

Code (MATLAB, Mathworks, Massachusetts, USA) was used to define the well depths based on the medium of choice and was used to define a computer-aided design program that controlled the operation of a laser system. The excimer machining laser used for this application was a 6 mJ, 200 Hz, 248 nm (Lasershot, Optec Inc., San Diego, CA USA and Frameries, Belgium) KrF laser machining system. A grid was defined using the method of quadratic residues described above and the well depth was calculated for each increment along the X and Y directions. The parameters needed for determining well depth are the speed of sound in glass, *c* = 4550 m/s, and the operating frequency of the transducer, *ω_r_* = 2*πf* where *f* in this case is 7 MHz, corresponding to the fundamental frequency for 500 *μ*m thick 128YX lithium niobate^34^. The well depth ranged up to 309 *μ*m and required between three and ten laser machining passes. The machined diffuser was bonded to the transducer face using a UV-curable epoxy (NOA81, Norland Products Inc, Cranbury, NJ USA) using previously described techniques^42^. This fabrication technique enables the miniaturization of devices that are capable of producing diffuse acoustic fields irrespective of the nature of the enclosed volume, as determined using surface and domain measurements as follows. A scanning laser Doppler vibrometer (UHF-120SV, Polytec, Waldbronn, Germany) was used to characterize the displacement of the substrate when actuated. Measurements of the pressure output from the transducer were performed using a fiber optic hydrophone (FOHS92, Precision Acoustics, Dorchester, UK).

### Ultrasound field simulation

Finite element analysis (COMSOL 5.5, Comsol Inc., Los Angeles, CA USA) was used to simulate the system as a linear media with a time-dependent acoustic pressure field present in two dimensions. The boundaries between the coupling fluid and the cover slip, and the cover slip and the media above it were defined as acoustic-structure boundaries, where there is fluid load on the structure due to pressure waves originating from the ultrasound source and structural acceleration on the fluid domain across the fluid-solid boundary. This results in stress build-up in the cover slip that is translated to the fluid domain above it for the duration of the stimulation. The coverslip was defined to have the elastic properties of silica glass, with an isotropic Young’s modulus of 73.1 GPa, a density of 2203 kg/m^3^ and a Poisson’s ratio of 0.17. The distance between the transducer and the upper boundary is 5 mm. The simulations were conducted in the time domain, with a 20 *μ*s burst followed by an 10 *μ*s dwell to observe changes in the pressure field both during and after the stimulus. The acoustic field was modeled as a sinusoidal input to the transducer, *d* = 5sin(*ωt*) nm, and the fluid domain was defined to have the properties of water (*ρ* = 1000 kg/m^3^, *c* = 1500 m/s). The maximum mesh size was chosen so that all element sizes were always less than one-eighth of a wavelength, and the data were exported every 0.05 *μ*s so that a frequency range of up to 10 MHz could be analyzed. The cover slip was defined to be 500 *μ*m thick and spanned the entire width of the domain. The spatial step chosen for plotting the isofrequency contours was less then *k*_max_ = *ωc*^−1^.

### Imaging rig for ultrasound stimulation

An upright epi-fluorescent microscope (Imager M2, Carl Zeiss GmbH, Gottingen, Germany) was used for the *in vitro* experiments. For this application we used our transducer assembly placed in a heated stage fixture set to 37°C underneath the cell chamber, which ensures homeostatis. Stimulus frequency and duration was controlled by a waveform generator (33600A Series, Keysight, CA USA), and the pressure was controlled through a 300-W amplifier (VTC2057574, Vox Technologies, Richardson, TX USA). Simultaneous calcium imaging was performed using a 40x water dipping objective at 16.6 frames per second with a camera (Orca Flash 4.0, Hamamatsu Photonics K.K., Japan) and a GFP filter.

### HEK293 cell culture and transfection

HEK293 cells (ATCC CRL-1573) were cultured using a standard procedure in DMEM supplemented with 10% fetal bovine serum (FBS) and 20 mM glutamine in a 37°C and 5% CO_2_ incubator. Cells beyond passage thirty were discarded and a new aliquot was thawed. A stable calcium reporter line was generated with a GCaMP6f lentivirus (Cellomics Technology PLV-10181-50) followed by fluorescence-activated cell sorting (FACS). For diffuser experiments, GCaMP6f-expressing HEK cells were seeded on a twelve-well cell culture plates with 18-mm glass coverslips coated with poly-D-lysine (PDL) (10 *μ*g/*μ*l; P6407, Sigma-Aldrich, St. Louis, Missouri, USA) for 1-2h. Coverslips were washed with (Milli-Q) ultrapure water and cells were seeded at a density of 250000 cells/well. Cells were transfected with lipofectamine LTX Reagent (15338100, ThermoFisher Scientific, Massachusetts, USA) according to the manufacturer’s protocol and 24 hours after plating, using 500 ng DNA of the clone of interest for each well. Cells were kept at 37°C for an additional 24 h before imaging on our ultrasound stimulation setup. For imaging, cover slips were mounted on a specialized chamber featuring an ultrasound transducer approximately 5 mm below the cover slip and a 10 mL reservoir of media above the cover slip. Once cells were in focus, an ultrasound pulse of 100 ms duration was delivered as described in previous sections while imaging with a 40X immersion objective, and a cell membrane profile was reconstructed and analyzed from these images (ImageJ, National Institutes of Health, Bethesda, Maryland, USA).

### Rat primary neuronal culture

Rat primary neuronal cultures were prepared from rat pup tissue at embryonic day (E) 18 containing combined cortex, hippocampus and ventricular zones. The tissue was obtained from BrainBits (catalog #: SDE-HCV) in Hibernate-E media and used the same day for dissociation following their protocol. Briefly, tissue was incubated in a solution of papain (BrainBits PAP) at 2 mg/mL for 30 min at 37°C and dissociated in Hibernate-E for one minute using one sterile 9” silanized Pasteur pipette with a fire-polished tip. The cell dispersion solution was centrifuged at 1100 rpm for 1 min, and the pellet was resuspended with 1 mL NbActiv1 (BrainBits NbActiv1 500 mL). The cell concentration was determined using a haemocytometer and neurons were plated in 12-well culture plates with 18-mm PDL-coated cover slips (Neuvitro Corporation GG-18-PDL) at a concentration of 900,000 cells/well. Neurons were then incubated at 37°C, 5% CO_2_, performing half media changes every 3-4 days with fresh NbActiv1 supplemented with PrimocinTM (InvivoGen ant-pm-1). For calcium imaging experiments, cells were infected with AAV9-hSyn-GCaMP6f (Addgene #100837-AAV9) at day 3 *in vitro* (DIV3) and a half media change was performed the next day. Neurons infected with GCaMP6f as stated above were infected with AAV9-hSyn-Cre (Addgene #105553-AAV9) and AAV9-hSyn-TRPA1-myc-DIO (Salk GT3 core) at DIV4 and a half media change was performed the next day. Cultures were incubated at 37°C, 5% CO_2_ until DIV10-12 and then imaged using the same equipment as for HEK cell experiments.

### Calcium imaging

Calcium imaging analysis was performed using custom scripts written as ImageJ macros. Transfected cells were segmented and cell fluorescence over time in the GCaMP6f channel was measured and stored in comma-delimited text (csv) files. Calcium data was analyzed using custom Python scripts. Calcium signals were normalized as ΔF/F using a 6 s baseline for each region of interest (ROI) and a peak detection algorithm with a fixed threshold of 0.25 was used to identify responsive cells after ultrasound stimulation.

### Ex-vivo hydrophone measurements

Hydrophone measurements were performed with a fiber-optic hydrophone (FOHS92, Precision Acoustics, Dorchester, UK) *ex vivo*. C57BL/6 mice (JAX 000664), aged 10-14 weeks, were sacrificed and decapitated. The skin over the skull was removed, followed by removal of the lower mandible, soft palate and hard palate. Once the ventral part of the brain was exposed, the mouse head preparation was placed dorsal side down on the diffuser assembly coupled with ultrasound gel, and the hydrophone tip was lowered into the ventral portion of the brain using a micromanipulator (Fig. S3). Animals used in this trial were group housed in an American Association for the Accreditation of Laboratory Animal Care approved vivarium on a 12 hour light/dark cycle, and all protocols were approved by the Institutional Animal Care and Use Committee of the Salk Institute for Biological Studies. Food and water were provided ad libitum, and nesting material was provided as enrichment. Mice were euthanized using CO_2_ according to approved protocols before decapitation and dissection.

## Acknowledgements

The authors are grateful to the University of California San Diego for provision of funds and facilities in support of this work. J.F. is grateful for funding for this work from the W.M. Keck Foundation via a SERF grant and, with S.C., from the National Institutes of Health via R01NS115591. S. C. is also grateful to the NIH in support of this work via grant R01MH MH111534. This work was performed at the Medically Advanced Devices Laboratory at the University of California, San Diego and the Chalasani lab at the Salk Institute of Biological Sciences. Fabrication was performed in part at the San Diego Nanotechnology Infrastructure (SDNI) of UCSD, a member of the National Nanotechnology Coordinated Infrastructure, which is supported by the National Science Foundation (Grant ECCS–1542148). The authors are grateful to John Roy and team in San Diego from Optec Laser Systems, for substantial training, assistance, and advice in laser machining throughout this effort. The authors would like to thank members of the Medically Advanced Devices Laboratory and the Chalasani Laboratory for feedback on this manuscript.

## Author contributions

J.F. and A.V. designed the research. A.V. built, characterized the devices and conducted experiments. A.V., FA., and N.B. analyzed data. M.D. and S.H.C. performed in-vitro studies. A.V. and U.M. performed the ex-vivo study. A.V. and J.F. wrote the paper.

